# Functional Balance between TCF21-Slug defines phenotypic plasticity and sub-classes in high-grade serous ovarian cancer

**DOI:** 10.1101/307934

**Authors:** Sagar S Varankar, Swapnil C Kamble, Avinash M Mali, Madhuri M More, Ancy Abraham, Brijesh Kumar, Kshama J. Pansare, Nivedhitha J Narayanan, Arijit Sen, Rahul D Dhake, Aparna N Joshi, Divya Midha, Mohit Kumar Jolly, Ying Dong, Judith A Clements, Sharmila A Bapat

**Author notes:** These authors contributed equally to this work (Sagar S Varankar, Swapnil C Kamble).

## Abstract

Cellular plasticity and transitional phenotypes add to complexities of cancer metastasis initiated by single cell epithelial to mesenchymal transition or cooperative cell migration (CCM). We identified novel regulatory cross-talks between Tcf21 and Slug in mediating phenotypic and migration plasticity in high-grade serous ovarian adenocarcinoma. Live imaging discerned CCM as being achieved either through rapid cell proliferation or sheet migration. Transitional states were enriched over the rigid epithelial or mesenchymal phenotypes under conditions of environmental stresses. The Tcf21-Slug interplay identified in HGSC tumors through effective stratification of subtypes also contributed to class-switching in response to disease progression or therapy. Our study effectively provides a framework for understanding the relevance of cellular plasticity in situ as a function of two transcription factors.

## Introduction

Intrinsic cellular identities originate during embryonic commitment and are fated for tissue specific spatio-temporal roles through cross-talks with niche components^1^. Destabilization of homeostasis alters the tissue milieu to revoke inherent cellular plasticity and restore primary phenotype(s), architecture and functions^2,3^. Processes such as the cooperative action of Slug and Sox9 for maintenance of basal and luminal epithelial cells in the mammary gland are diverted to define different cell fates in an aberrant context like cell transformation^4,5^. Innate capabilities and perturbed microenvironment thus align as distinct etiologies in cancer^6,7^. Different subtypes identified in epithelial ovarian carcinomas (EOC) as Stem-A, Stem-B, Epi-A, Epi-B and Mes correspond with epithelial (E), intermediate epithelial (iE), intermediate mesenchymal (iM) and mesenchymal (M) features^8,9^; a subsequent study reported iE and iM as an epithelial-mesenchymal hybrid (E-M)^10^. Bunching of disparate EOC misrepresents associations between subtypes and phenotypic states, especially since systematic understanding of the most aggressive EOC *viz*. heterogeneous high-grade serous adenocarcinomas (HGSC) has recently revealed diverse molecular pathways, mutations and gene expressions that correlate with distinct epithelial/differentiated (E/D), mesenchymal (M), proliferative (P), immunoreactive (IR) features ^11–15^.

Our previous analyses of HGSC expression datasets in The Cancer Genome Atlas (TCGA) resolved three molecular sub-classes associated with discrete mechanisms of metastases. One of these exhibited co-operative cell migration (CCM), another undergoes epithelialmesenchymal transition (EMT) while Class3 presents with mixed features^12,16^. Correlating these with other studies using the same datasets suggested CCM-Class tumors to present as P/D/iE sub-class, EMT-Class tumors as M/IR/iM while the heterogeneous Class3 tumors possibly represent IR/D phenotypes. In the present study we discern different HGSC cell phenotypes (E/M/intermediates) as a function of Tcf21 and Slug expression, frequency, localization and activity. Strikingly, epithelial features involve transcriptional repression of Slug by Tcf21. Functional evaluation associated different phenotypes with either EMT or CCM; the latter in turn was revealed as being either passive (pCCM, manifested as rapid proliferation to achieve cell displacement) or active (aCCM, sheet migration). Intermediate states altered Tcf21-Slug sub-cellular localization, exhibited transitions between EMT and aCCM (but not pCCM) and greater plasticity in response to positive/negative growth regulators. *In situ* assessment of the different phenotypes through an immuno-histochemistry-(IHC) and HC-based scoring of six markers in HGSC tumors not only reflected the CCM, EMT and double positive (DP-intermediate state) classes, but also revealed class switching following metastasis and/or therapeutic intervention. In summary, cellular phenotypes defined in terms of transcription factor (TF) mediated cellular plasticity associated with distinct migratory capabilities, may be relevant to the clinical presentation and management of HGSC.

## Results

### Differential Tcf21 and Slug expression defines phenotypic plasticity and modes of migration that are perturbed by environmental cues

Data from our previous study^16^ identified class-specific association of thirteen transcription factors (TFs; Supplementary Table1); AIF1,ETV7 were excluded from further analysis due to experimental complexities of assessing immunomodulation. Promoter occupancies between the remaining TFs derived from known consensus DNA binding sequences (Figs.1a,S1a) predicted co-regulation of *Tcf21,Gata4,Peg3,Gli1* and *Foxl2* that correlated positively with each other and negatively with *Snai2* (Slug) in the TCGA dataset (Fig.S1b). Profiling these TFs in class-representative cell lines robustly associated CCM-Class (OVCAR3 cells) with Tcf21 and EMT-Class (A4 cells) with Slug - TCF2^lo^ expression (Figs.1b,S1c). We further derived knockout (K/O) clones of the dominant TF followed by over-expression (OE) of the other TF in the 2 cell lines (Fig.S1d). OVCAR3 cells expressed high Tcf21, E-cadherin (*Cdh1*); Tcf21^K/O^(E2) exhibited high nuclear Slug, low *Cdh1*; A4 cells expressed high Slug, Vimentin (*Vim*) with low *Cdh1*; sustained expression of other EMT-TFs (Zeb1/2,Twist1,Snail,Prrx1) in the Slug^K/O^ (D7) led to minimally altered *Cdh1* and *Vim* despite nuclear Tcf21 (Fig.S1e); A4_Slug^K/O^Tcf21^OE^ (TCF_OE) lowered EMT-TFs to enhance *Cdh1* expression (Figs.1c,1d). High luciferase activity in OVCAR3 and TCF_OE cells (reporter assay for *Cdh1* promoter harboring Slug binding E-box sequences), and reduced activity in E2, A4 and D7 cells suggests phenotypic variability as a function of Tcf21 and Slug expression (Fig.S1f).

**Fig.1.**
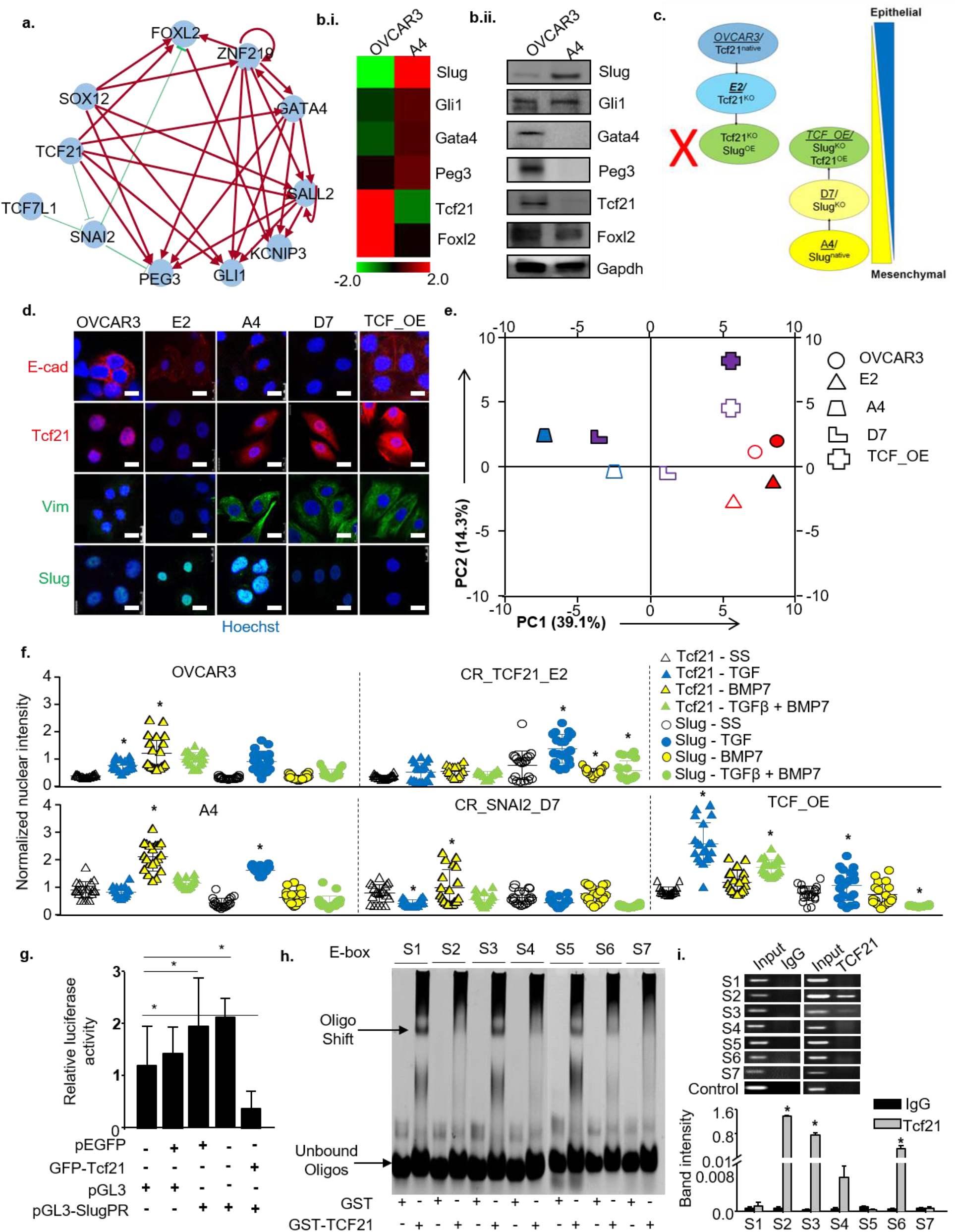
Tcf21 and Slug govern cellular phenotypes in HGSC. a. Predicted transcription factor (TF) network of cross-regulatory mechanisms, red arrows and green bars represent likely activator or repressor functions; b. Expression profiling of *Snai2*(*Slug*),*Gli1,Gata4, Peg3,Tcf21,Foxl2* in OVCAR3 and A4 cells by (i) qPCR (heatmap) and (ii) immunoblotting, GAPDH used as endogenous control; c. Flow-chart depicting step-wise generation of varied Tcf21 and Slug derivative clones from parental cells OVCAR3 and A4, the color gradient depicts an epithelial–mesenchymal (E-M) axis along which parental and derivative cells are placed; d. Immunostaining of Tcf21,Slug,Ecad,Vim in parental and derivative cells, nuclei visualized with Hoechst, Scale bars-20μm; e. Principle component (PC) analysis of time-lapse imaging-based migration data of parental and derivative cells, PC1 – variance between displacement (Final Y) and velocity *vs*. nearest neighbours, PC2 – variance between displacement and velocity, color gradient depicts migratory modes, blue-EMT: purple-aCCM: red-pCCM, filled and empty shapes indicate presence and absence of serum respectively; f. Representative graphs depicting frequency and intensity of Tcf21 and Slug expressing nuclei in parental and derivative cells, data depicts fluorescence intensities measured across five microscopic fields; g. Reporter assays to determine effects of Tcf21 on wild type Slug promoter in A4 cells, luciferase activity normalized to untransfected cells, pEGFP:empty GFP vector, GFP-Tcf21:Tcf21-pEGFPC1 recombinant vector, pGL3:native luciferase vector, pGL3-SlugPR:pGL3 containing Slug promoter as defined in Fig.S3; h. Representative EMSA of GST–Tcf21 recombinant protein binding to E-boxes in Slug promoter region in a cell free system, GST protein used as control; i. ChIP-PCR analysis of Tcf21 binding to E-box elements in Slug promoter in OVCAR3 cells, IgG used as isotype control, cross-linked fragmented DNA considered as Input. All data are representative of experiments performed in triplicate and are depicted as mean ± SEM. *p<0.05, **p<0.01, ***p<0.001.

Further assessment of biological functions (adherent colony, suspended spheroid formation, migration, invasion) revealed only the rates and modes of migration to be differential between parental and derivative clones, and were positively influenced by serum (Fig.S1g; Supplementary movies 1 – 2). OVCAR3 and E2 cells exhibit proliferation-driven cell displacement that we termed passive CCM (pCCM), A4 mediated rapid wound healing through a combination of single cell and active sheet migration (EMT, aCCM) while its derivatives D7, TCF_OE exhibit aCCM. Principal component (PC) analyses of time-lapse cell imaging using quantifiable metrics of cell displacement (Final Y), velocity and nearest neighbors supports differential migration (Fig.1e). PC1 (variance between velocity and displacement *vs*. number of nearest neighbors) confirmed EMT-aCCM based migration in A4 (low nearest neighbors, high velocity), aCCM in D7 and TCF_OE (high nearest neighbors, lower velocity) and sustained cell proliferation driven by pCCM in E2 and OVCAR3 cells (highest nearest neighbors, lowest velocity). PC2 (variance between displacement *vs*. velocity) was minimum in A4, D7, OVCAR3 and maximum in TCF_OE and E2 cells. Differential nearest neighbors between A4 and D7 appeared to be a useful metric in differentiating EMT from aCCM apart from the visualization in real time imaging. These data of a gradient of velocities, nearest neighbors and displacement effectively define an E to M spectrum as shown in Fig.1c.

We evaluated effects of TGFβ and BMP7 that are involved in normal ovulation, on the generated spectrum. OVCAR3 cells express low levels of BMPR2 and TGFβR2 receptors and are refractory to both growth factors; E2 (BMPR2^lo^/TGFβR2^mod^) responds poorly to BMP7, while TGFβ induced nuclear Slug expression (Figs.1f,S2a,S2b,S2c). A4, D7, TCF_OE (BMPR2^mod^) respond to BMP7 through nuclear localization of Tcf21, enhanced Cdh1 expression, reduced Slug and wound healing (Fig.S2d). TGFβ represses Tcf21, Cdh1 in A4 and D7 cells (TGFβR2^hi^) concomitantly with increased Slug, Vim expression and enhanced migration, while TCF_OE (TGFβR2^lo^) responds poorly to TGFβ treatment. Cell responses to combination of these two growth factors agreed with the respective receptor status; Vim levels were not significantly altered by either growth factor. PC analyses of time-lapse cell imaging affirmed EMT in A4 in the presence of TGFβ, pCCM in E2 and OVCAR3, aCCM in TCF_OE, while D7 cells shifted from aCCM towards EMT (Fig.S2e; Supplementary Movie 3; PC1 and PC2 as defined earlier). These results reveal a complexity wherein E and M states may not undergo a simple binary phenotypic switch, but rely on cooperation between intrinsic TF expression and receptor status to mediate migration in response to environmental stimuli.

### TCF21 is a transcriptional repressor of Slug

The next step was to probe possible cross-regulation between TCF21 and Slug. Luciferase activity (reporter assay for *SNAI2* promoter harboring TCF21 consensus E-box sequences) was severely reduced in A4 cells expressing Tcf21 (Figs.S3,1g). *In vitro* binding assays affirmed physical interactions between recombinant Tcf21 and S1,S3,S5 *SNAI2* E-boxes, while probing of Tcf21-bound chromatin complexes in OVCAR3 cells through immunoprecipitation indicated affinity for S2,S3 *SNAI2* E-boxes (Figs. 1h,1i). This suggests novel regulation of *SNAI2* at the transcriptional level by Tcf21. Similar regulation of TCF21 expression by Slug was not observed.

### HGSC tumors and cell lines intrinsically display similar spectrum of phenotypes and modes of migration

Correlations between several phenotypic markers in TCGA expression data generated four clusters (Fig S4a). C1 comprised of epithelial cell junctions and cytoskeletal molecules, C2 genes reportedly associate with intermediary phenotypes in several tissues^17–19^, C3 represents cell junction components with mesenchymal features^11^, while C4 included intermediate filaments, endothelial and stromal components^20,21^. Hierarchical clustering of epithelial and mesenchymal TFs in this data yielded five distinct tumor groups (Fig S4b). Two of these were exclusively E and M, a third was considered E-M by virtue of a double-positive signature; a group identified as iE expressed E markers, Snail, Twist1 but lacked Tcf21, while the last tumor group with M features, yet low EMT-TF expression was assigned as iM. Parallel screening of E, M markers and TF transcripts across a panel of HGSC cell lines resolved phenotypic states through definitive correlations between E (Cdh1,Ck19), M (Vim,FAP) markers, Tcf21, Slug and Prrx1 isoforms 1a and 1b^22^ (Figs.S4c,S4d,S4e; Prrx1a, Prrx1b transcripts - P1a, P1b respectively, protein represented as Prrx1; non-availability of isoform-specific Prrx1 antibodies restricted further definitive studies). OVCAR3, OVCA432 (high E, nuclear Tcf21, low P1a-P1b-nuclear Prrx1, low nuclear Slug) and OVMZ6 cells with an inverse profile defined the E and M ends of the spectrum; CAOV3 and OV90 were identified as iE (high E, FAP, nuclear Tcf21, low P1a-P1b-moderate nuclear Prrx1, low nuclear Slug); OVCA420 and PEO14 were E-M (E, M markers, all TFs in nucleus), while A4 represents iM (CK19, M markers, low cytoplasmic Tcf21, high P1b, nuclear Prrx1 and Slug). These phenotypes could be perturbed by external stress such as serum deprivation that increases Cdh1 and nuclear Tcf21, concomitantly decreasing Slug, FAP and Vim to shift the spectrum towards the E end (Figs.2a,2b).

**Fig.2.**
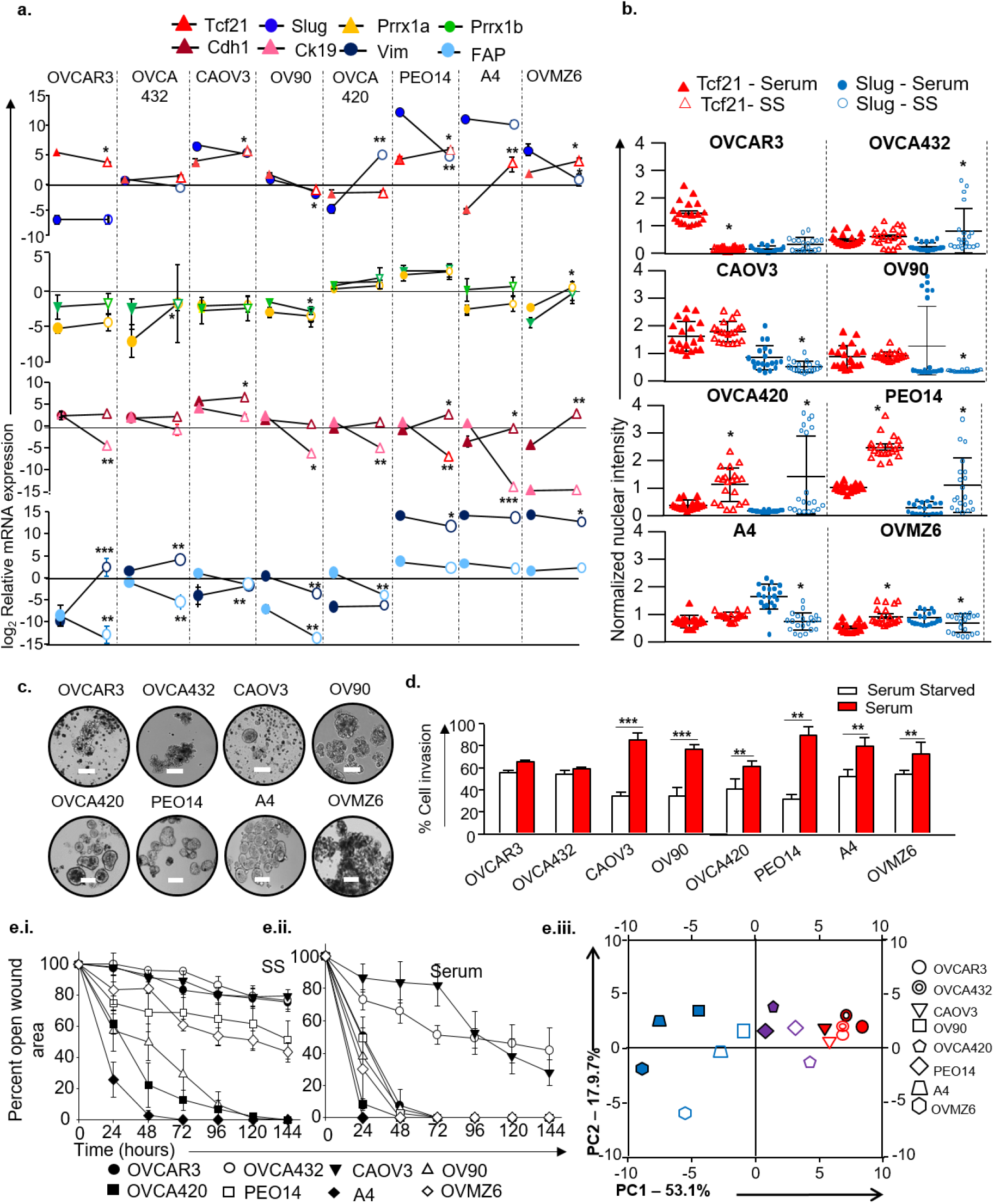
Resolution of the epithelial – mesenchymal phenotypic axis highlights differential functionalities as a function of Tcf21 and Slug localization. a. Expression profiles of Tcf21, Slug, Prrx1 isoforms, Cdh1, Ck19, Vim and FAP in HGSC cell lines at steady states and post serum deprivation normalized with GAPDH. Filled shapes indicate the presence of serum and empty shapes indicate that cells were serum starved; b. Representative graphs depicting frequency and intensity of Tcf21 and Slug positive nuclei in HGSC cell lines. Data depicts fluorescence intensities measured across five microscopic fields; c. Representative images of suspension culture morphology (Scale bars-200μ); d. Percent cell invasion in Boyden chamber matrigel assay in the absence and presence of serum; e. Efficacy of wound healing in the (i) absence and (ii) presence of serum. e.iii. Principle component analysis used to project segregation of migratory modes in the phenotypic spectrum based on quantitative metrics - Final ‘Y’, velocity and nearest neighbors - emerging from time-lapse based migration data. Color gradient represents the migratory modes of each cell line. Filled shapes indicate the presence of serum and empty shapes indicate that cells were serum starved; All data are representative of experiments performed in triplicate and are depicted as mean + SEM, *p<0.05, **p<0.01, ***p<0.001.

Probing phenotypic - functional associations indicated intermediate states could generate organized spheroids in suspension while E and M cells form clumps/clusters (Fig.2c). Except the E state, all cell lines were highly invasive in response to a serum gradient (Fig.2d). Differential migration capabilities were evident wherein A4, OV90 and OVCA420 cells filled the open wound area in absence of serum as opposed to E/iE and surprisingly M cells that required serum to initiate migration (Fig.2e.i,2e.ii). Live imaging and PC analyses revealed pCCM in E and iE (CAOV3), aCCM in E-M, coupling of aCCM-EMT in iM and EMT in M cells (Supplementary Movies 4 – 5; Fig 2e.iii). Interestingly, serum deprivation led to nuclear localization of Slug in a few E-M state cells that possibly induces the rare EMT events visualized in live imaging.

### Micro-environment influenced phenotypic inter-conversions are captured by nuclear Tcf21 and Slug expression

Cell line responses to BMP7 and TGFβ were regulated by expression of their respective receptors (Fig S5a). E and iE (CAOV3) cells exhibit low to moderate levels of both receptors, were least responsive and maintained a rigid epithelial phenotype. E-M, iM and M cell lines exhibited high receptor expression, hence while BMP7 induced Cdh1, nuclear Tcf21 and reduced Slug levels, TGFβ exposure up-regulated nuclear Slug concomitantly with reduced Cdh1 and nuclear Tcf21 (Figs 3a,S5b,S5c). Dominant effects of BMP7 over TGFβ were observed in E-M and iM cell lines (high Cdh1, nuclear Tcf21, low Slug) when the growth factors were provided in combination. PC analysis of expression profiles indicated minimal variance as a function of TFs along PC1 in OVCAR3, OVCA432 (E), CAOV3 (iE) and A4 (iM) cells; moderate in PEO14 (E-M) and OVMZ6 (M) cells and maximum variance in OV90 (iE) OVCA420 (E-M;Figs.3b,S5b). PC2 as a function of Cdh1 and Vim strengthened these observations (Figs 3b,S5b). Correspondingly, rate of migration is restricted by BMP7 and enhanced by TGFβ in E-M, iM and M, while minimal changes were evident in E and iE cells (Fig S5d). Together, this affirms phenotypic rigidity of E cells while reflecting on cellular plasticity of the others.

**Fig3.**
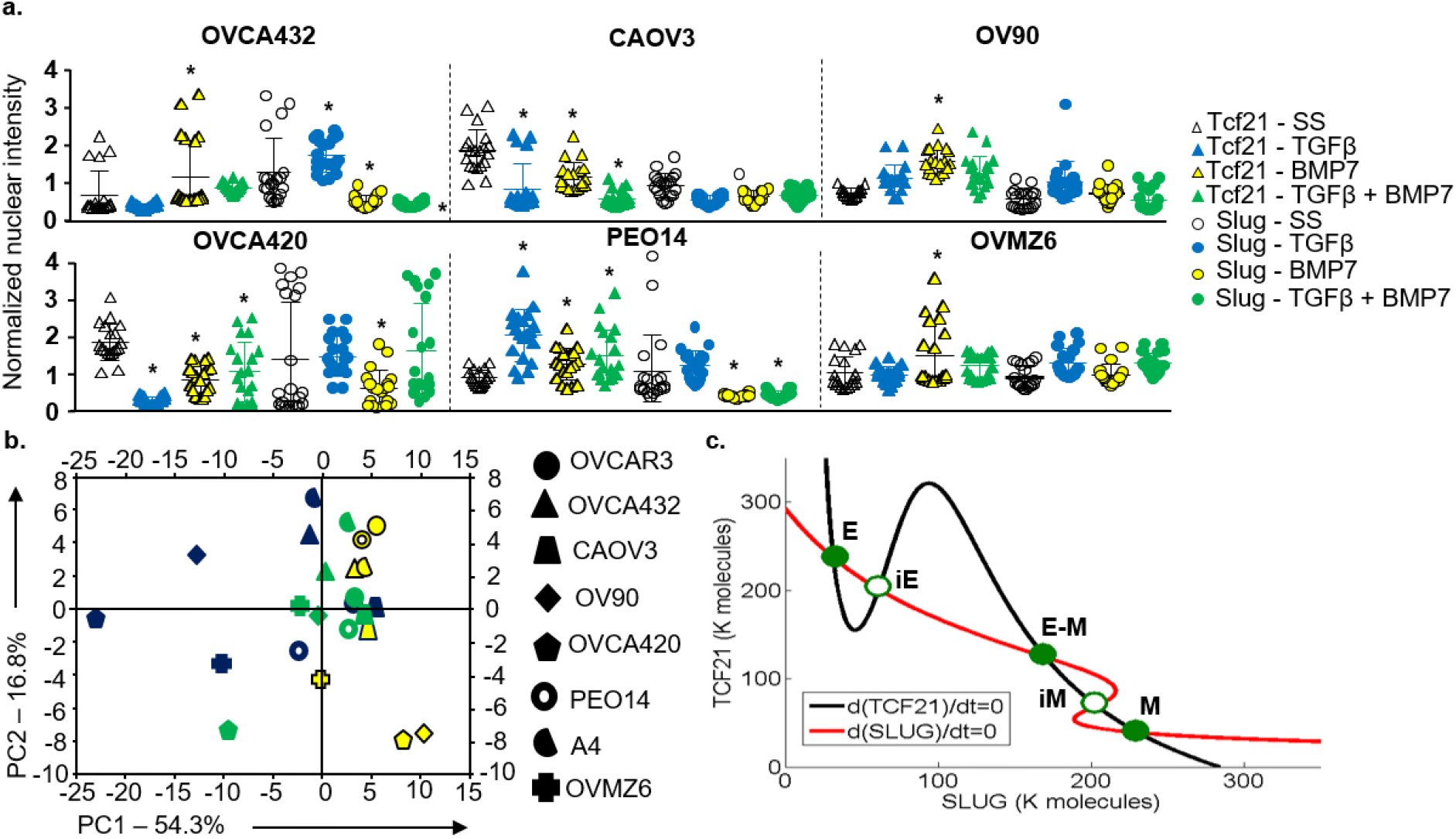
Phenotypic axis highlights differential responsiveness to micro-environmental regulation as a function of Tcf21 and Slug localization. a. Representative graphs depicting frequency and intensity of Tcf21 and Slug positive nuclei in HGSC cell lines exposed to BMP7, TGFβ and BMP7+TGFβ. Data depicts fluorescence intensities measured across five microscopic fields; b. Principle component analysis used to project phenotypic switches based on mRNA profiles following TGFβ and BMP7 exposure. Yellow, blue and green filled shapes correspond to cells exposed to BMP7, TGFβ and BMP7+TGFβ respectively; c. Mathematical model depicting variation in steady-state levels of TCF21 as a function of SLUG and vice-versa (nullclines shown as red and black curves), curve intersections mark stable cellular states indicated by solid green circles (TCF21^high^-SLUG^low^, TCF21^low^-SLUG^high^ and Tcf21^mod^-Slug^mod^), while hollow circles indicate metastable cellular states that correspond to different phenotypes observed along the E-M spectrum; all data are representative of experiments performed at least in triplicate.

Integrating Tcf21 and Slug transcript profiles across all cell lines and derivatives under varying culture conditions (steady-state, serum-deprivation, growth factor exposure) with TF-promoter binding studies aided the generation of a mathematical model that identified three stable phenotypes *viz*. TCF21^high^-SLUG^low^ (E), TCF21^low^-SLUG^high^ (M) and Tcf21^mod^-Slug^mod^ (E-M); iE and iM were depicted as metastable states during phenotype switching wherein drastic fluctuations in Tcf21 and Slug expression correlated with transitions at the E and M ends respectively (Fig.3c). Importantly, functional assessment assigns cellular rigidity to E and M states and plasticity to intermediate states, which assigns definitive significance of intrinsic phenotypes in biological systems. Conclusively, TCF21-Slug expression and sub-cellular localization correlates with cellular states across a spectrum and are possible determinants of functionality in migration and phenotypic plasticity.

### Scoring of Phenotype-Specific Biomarkers in HGSC Xenografts and TMAs

We further established immunohistochemistry (IHC) and histochemistry (HC) based read-outs in xenografts of a panel of six markers including Tcf21 and Slug to explore correlations between the *in vitro* phenotypes and HGSC molecular classes. Based on our earlier molecular stratification, additional markers included Cdh1, PARP1 (CCM/E class), Slug, hyaluronan (HA), and AnnexinA2 (ANXA2; EMT/M class; Fig S6a). An inability to correlate variations in Vim protein expression with TF expression or growth factor stimuli refrained its inclusion. Reproducible standard operating protocols (SOPs) were established (available on request) to address pre-analytic, analytic, and post-analytic parameters. Derivation of universal guidelines for marker scoring based on the metrics of marker frequency, intensity and localization (Fig S6b), and development of a reference score sheet using appropriate healthy human tissues based on the Human Protein Atlas (HPA)^23^ was further undertaken (Fig S6c). Subjectivity of analyses was minimized by collecting blinded scores from 5 observers (SCK,AS,RDD,ANJ,DM) followed by a comprehensive pathology review to arrive at a consensus in case of difference in opinions.

Tcf21 expression in xenograft sections was either dominantly nuclear (CAOV3,PEO14), cytoplasmic (OVMZ6, OVCAR3) or negligible (A4); similarly Slug was nuclear (OVMZ6, A4), cytoplasmic (CAOV3) or absent (OVCAR3,PEO14;Fig.4a.i). Moderate intensity of Cdh1 at the cell membrane was observed in 50% of CAOV3 and OVCAR3 xenografts, but was lower in OVMZ6, A4 and PEO14. Significant, high intensity expression of nuclear PARP1 was evident only in OVCAR3 xenografts, while only OVMZ6 and A4 xenografts exhibited ANXA2 expression. High frequency, moderate intensity of HA was observed in CAOV3 and A4, low to moderate frequency and weak intensity in OVCAR3 and PEO14 xenografts. Consensus marker scores consolidated by the pathologist panel for each marker and xenograft (S_Freq_, S_Int_, S_Loc_;Table 1) were used to derive specific Biomarker Indices (BI; equation i). Class-indices representing class-specific metrics of consolidated markers expression were derived from class-specific BI (CI_EMT_ and CI_CCM_; equations ii & iii; Table 1). On applying the distribution of median CI_EMT_ *vs*. CI_CCM_ values in class stratification, OVCAR3 xenografts were identified as CCM-class, A4 and OVMZ6 as EMT-class and CAOV3 and PEO14 as double positive (DP;Fig.4a.ii.). Importantly, this scoring achieved inclusiveness of the mixed/heterogeneous Class3 tumors by relative association of marker expression as CI_EMT_ and CI_CCM_ to correlate with the phenotypic plasticity observed *in vitro* (Fig.3a).

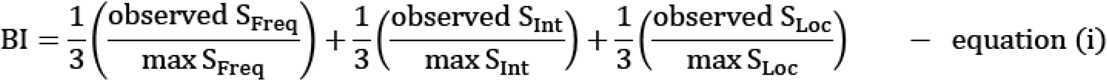

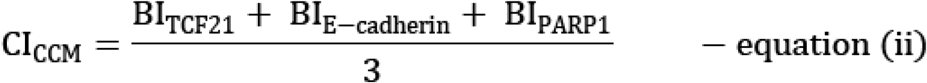

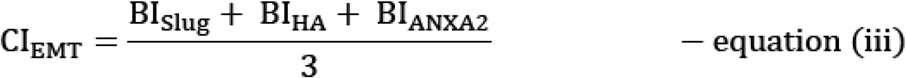

**Fig4.**
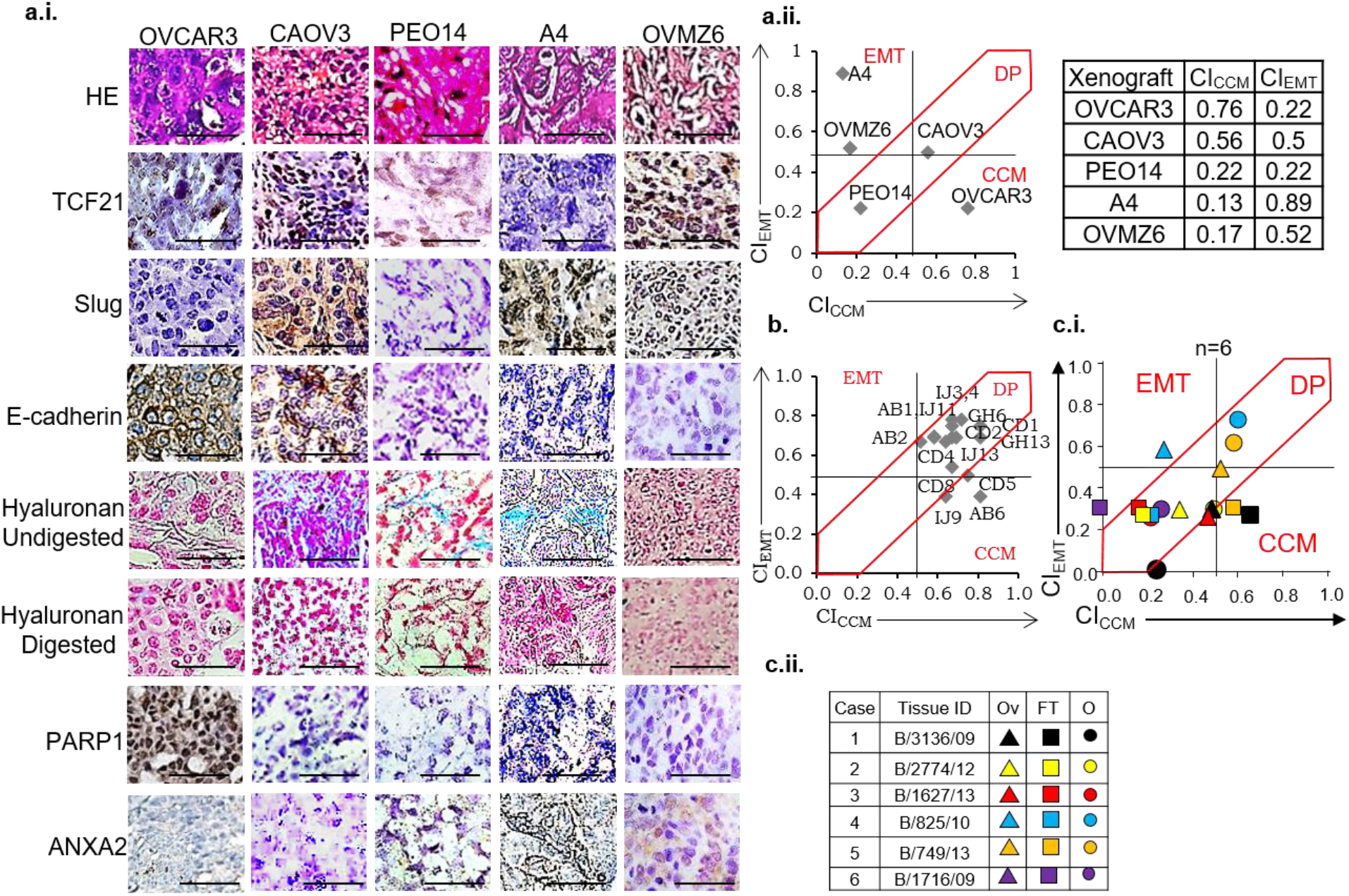
Class associations and marker scoring guidelines. a.i. Representative images of HGSC xenograft stained section for: Row 1 - H&E (hematoxylin and eosin), Rows 2,3,4,7,8 represent IHC-based identification of TCF21, Slug, Ecadherin, PARP1 and ANXA2, Rows 5 & 6 represent HC-based identification of HA fibers in untreated and hyaluronidase digested sections respectively, Scale bars-50μm; a.ii. Scatter plots of CI_CCM_ *vs*. CI_EMT_ derived from xenograft scoring; b. Similar scoring of tissue microarrays; c.i. Scoring of HGSC chemo-naive cases - tumors detected in ovary (Ov), fallopian tube (FT) and omentum (O); c.ii. Reference case-chart for c.i.

**Table1.** Scores, Biomarker and Class Indices (BI and CI respectively) for CCM and EMT markers in xenografts

**a. CCM Markers**
| Xenograft | TCF21 | | | | E-cadherin | | | | PARP1 | | | | $CI_E$ |
| --- | --- | --- | --- | --- | --- | --- | --- | --- | --- | --- | --- | --- | --- |
| | $S_{Freq}$ | $S_{Int}$ | $S_{Loc}$ | $BI_{TCF21}$ | $S_{Freq}$ | $S_{Int}$ | $S_{Loc}$ | $BI_{CDH1}$ | $S_{Freq}$ | $S_{Int}$ | $S_{Loc}$ | $BI_{PARP1}$ | |
| CAOV3 | 2 | 2 | 2 | 0.78 | 3 | 2 | 2 | 0.89 | 0 | 0 | 0 | 0 | 0.56 |
| OVMZ6 | 2 | 1 | 1 | 0.5 | 0 | 0 | 0 | 0 | 0 | 0 | 0 | 0 | 0.17 |
| A4 | 1 | 1 | 1 | 0.39 | 0 | 0 | 0 | 0 | 0 | 0 | 0 | 0 | 0.13 |
| OVCAR3 | 2 | 3 | 1 | 0.72 | 2 | 2 | 2 | 0.78 | 2 | 2 | 2 | 0.78 | 0.76 |
| PEO14 | 1 | 2 | 2 | 0.67 | 0 | 0 | 0 | 0 | 0 | 0 | 0 | 0 | 0.22 |

**b. EMT Markers**
| Xenograft | Slug | | | | HA | | | | ANXA2 | | | | $CI_M$ |
| --- | --- | --- | --- | --- | --- | --- | --- | --- | --- | --- | --- | --- | --- |
| | $S_{Freq}$ | $S_{Int}$ | $S_{Loc}$ | $BI_{Slug}$ | $S_{Freq}$ | $S_{Int}$ | $S_{Loc}$ | $BI_{HA}$ | $S_{Freq}$ | $S_{Int}$ | $S_{Loc}$ | $BI_{ANXA2}$ | |
| CAOV3 | 3 | 1 | 1 | 0.61 | 3 | 2 | 2 | 0.89 | 0 | 0 | 0 | 0 | 0.5 |
| OVMZ6 | 2 | 1 | 2 | 0.67 | 0 | 0 | 0 | 0 | 3 | 2 | 2 | 0.89 | 0.52 |
| A4 | 3 | 3 | 2 | 1 | 3 | 2 | 2 | 0.89 | 3 | 1 | 2 | 0.78 | 0.89 |
| OVCAR3 | 0 | 0 | 0 | 0 | 2 | 1 | 2 | 0.67 | 0 | 0 | 0 | 0 | 0.22 |
| PEO14 | 0 | 0 | 0 | 0 | 2 | 1 | 2 | 0.67 | 0 | 0 | 0 | 0 | 0.22 |

Scoring and tumor stratification guidelines were applied to commercial TMAs (duplicate cores per sample including 2 normal ovary + 13 HGSC cases among other ovarian cancer subtypes) restricted to four biomarkers (TCF21, Cdh1, Slug, HA; due to availability of limited consecutive slides) with appropriately modified equations (ii) and (iii;Supplementary Table 2). Most HGSC TMA-cores expressed high intensity cytoplasmic TCF21, moderate nuclear expression of which was present in ~5-10% of tumor cells. Except in four cases, Slug expression was weak to moderate cytoplasmic; nuclear localization was evident in 5-10% tumor cells. Moderate expression of Cdh1 was evident in all TMAs, but specifically at the cell membranes in 50% of the samples, while moderate intensity of extracellular HA fibers in tumor cell nests, was frequently observed. Consolidation of biomarker scores of each case, computation of BI and CI values followed by plotting distribution of CI_CCM_ *vs*. CI_EMT_ indicated 3 HGSC cores to represent CCM-Class while all remaining belonged to DP Class; EMT-Class was unrepresented (Fig.4b).

### Evaluation of clinical samples associates CCM-markers with metastases and chemotherapy

Towards assessing further clinical representation, we obtained and stratified 163 tumor samples pathologically diagnosed as HGSC from 96 patients. These included primary tumors [ovary (T), fallopian tube (FT)] and metastatic tumors [omentum (O), peritoneal ascites-derived cell blocks (A); Supplementary Table 3]. BI-CI scores and CI_CCM_-CI_EMT_ values for each tumor were computed (Supplementary Table 4) followed by evaluation of CI distribution and class assignment in the following groups –

**Group A.** Between group analyses of chemo-naïve (CN) tumors (T *vs*. FT *vs*. O;n=6),
**Group B.** Within group analyses of tumors at the above sites in CN cases (50-T, 7-FT, 26-O, 4-A) and chemo-treated (CT) cases (52-T, 2-FT, 17-O, 2-A),
**Group C.** Within group analyses of T-O pairs from either CN (n=17) or CT (n=16) cases, and
**Group D.** Between group analyses of tumor samples of the same case before and after therapy (T, A and/or O;n=6).

All three classes (CCM, EMT, DP) were represented in Group A. Primary tumors and metastases of case B/3136/09 predominantly expressed CCM-markers, while interestingly, metastases of all other cases segregated into DP class even when the primary tumor represented another class (Fig.4c). CN as well as CT T-tumors in Group B stratified into DP, CCM- or EMT- classes with increased representation of CCM-class following treatment (Figs.5a-i,5a-iii). CN FT-tumors also stratified into these three classes, while those after treatment were either CCM- or DP-Class (Figs.5a-i,5a-iii). O-tumor deposits were predominantly DP with marginally increased frequency of CCM- and EMT-markers following treatment (Figs.5a-ii,5a-iii); CN- as well CT-A cell blocks frequently presented as CCM-class (Figs.5a-ii,5a-iii). Overall, DP-class was better represented in ‘Group B’ CN-tumors (Supplementary Table 5) as compared to xenografts or TMAs (Table 1, Supplementary Table 2 respectively). Further comparison of the CI_CCM_ and CI_EMT_ means (M-CI_CCM_ and M-CI_EMT_ respectively) in evaluating effects of metastases and therapy (T, FT and O tumors from the same patient in CN *vs*. CT cases) revealed lower M-CI_CCM_ of CN: T-O tumors (Group C) than in CN: T-F-O tumors (Group B) while M-CI_CMM_ of CT: T-O tumors was enhanced (Fig.5b, Supplementary Table 6). Similarities between M-CI_EMT_ (CN: T-O *vs*. CN: T-F-O) and (CN: T-O *vs*. CT: T-O) tumors group suggested minimal effects of disease progression on expression of EMT-markers. Enrichment of CCM- and DP-subtypes in CT samples from Groups B and C could indicate either a predominance of such samples in our dataset, or therapy acquired marker expression. Further, tumor progression (T to O) identified metastases-associated class-switching in CN as well as CT tumors of Group C (Figs.5c-i,5c-ii); lack of directionality towards a specific phenotype suggests additional influences on marker expression. Within Group D (6 cases of T-O-A CN and CT tumors), Cases1 and 2 conformed to CCM-class after disease progression and treatment, while the remaining exhibited class-switching (Fig.5d). Cases 3 and 6 switched from a DP-profile of ovarian tumors to CCM-Class following treatment; Cases 4 and 5 was associated with heterogeneity of marker expression between different samples. These findings support metastasis / chemotherapy induced class-switching of HGSC tumors towards a CCM subtype.

**Fig5.**
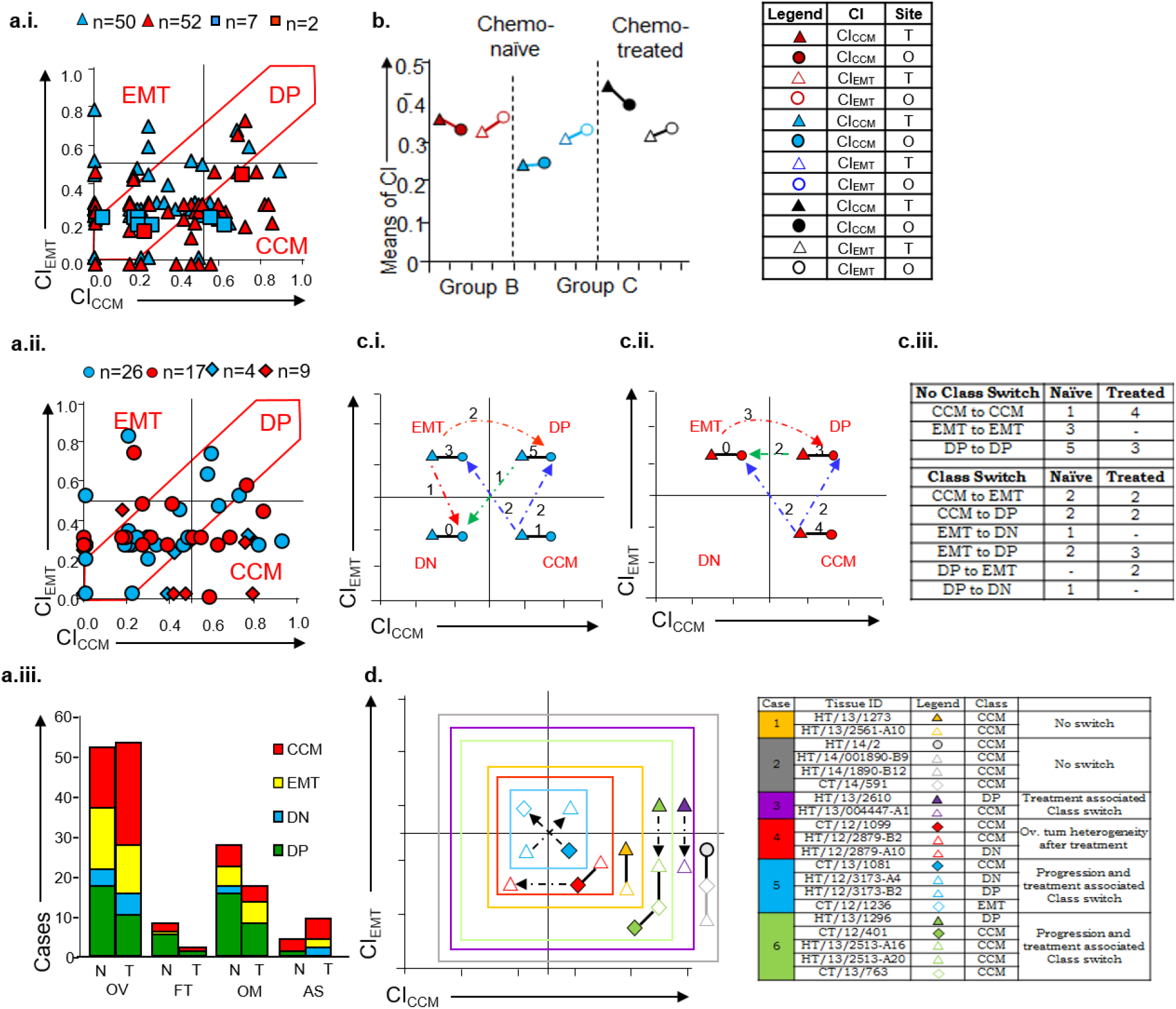
Chemotherapy induced alteration of the phenotypic spectrum. a. Scatter plots of CI_CCM_ *vs*. CI_EMT_ distribution for single chemo-naïve or -treated (red and blue shapes respectively) tumors from (i) primary tumors: ovarian Δ and fallopian tube □ and (ii) metastases: omentum ○ and ascites cell blocks ◊ a.iii. Graphical representation of Group A tumors (chemo-naïve - N; chemo-treated - T); b. Means plot comparing CI_CCM_ with CI_EMT_ of tumors from Groups B and C respectively; c. Ovarian and omental tumors tissues of i) Chemo-naïve (n=17) and ii) chemo-treated (n=16) patient cohort. d. Class switching in tumors collected from patients (n=6) prior to (filled shapes) and post chemo-therapy (empty shapes) as assessed in ascites (diamond), primary tumor (triangle) and omentum (circle); All data are representative of experiments performed in triplicate and are depicted as mean ± SEM. *p<0.05, **p<0.01, ***p<0.001.

### HGSC tumors at different sites exhibit molecular heterogeneity and class-switching

Class-switching in paired samples led us to examine for similar effects of paclitaxel in cell lines. PC analysis of live migration (PC1,PC2 as described earlier) identified a switch from EMT to aCCM mode for M and iM cells after treatment (maximum variance along both PCs), E-M and iE adhered to their characteristic modes of migration albeit at reduced velocities (moderate variance along both PCs) while pCCM phenotype in E cells was unaltered (Fig.6a, Supplementary Movies 6 – 7). These altered migratory metrics across the spectrum were associated with increased nuclear Tcf21 concurrently with reduced Slug expression (Fig.6b). Based on previous derivation of biomarker indices for clinical samples, expression profiles (intensity) and sub-cellular localization of Tcf21, Slug were utilized to derive TF indices as a means of capturing phenotypic switches *in vitro* (equation iv).

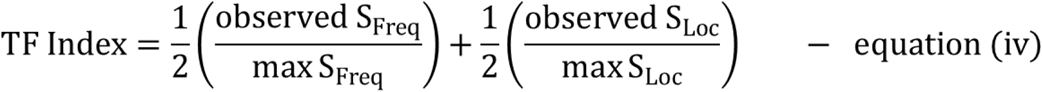

**Fig6.**
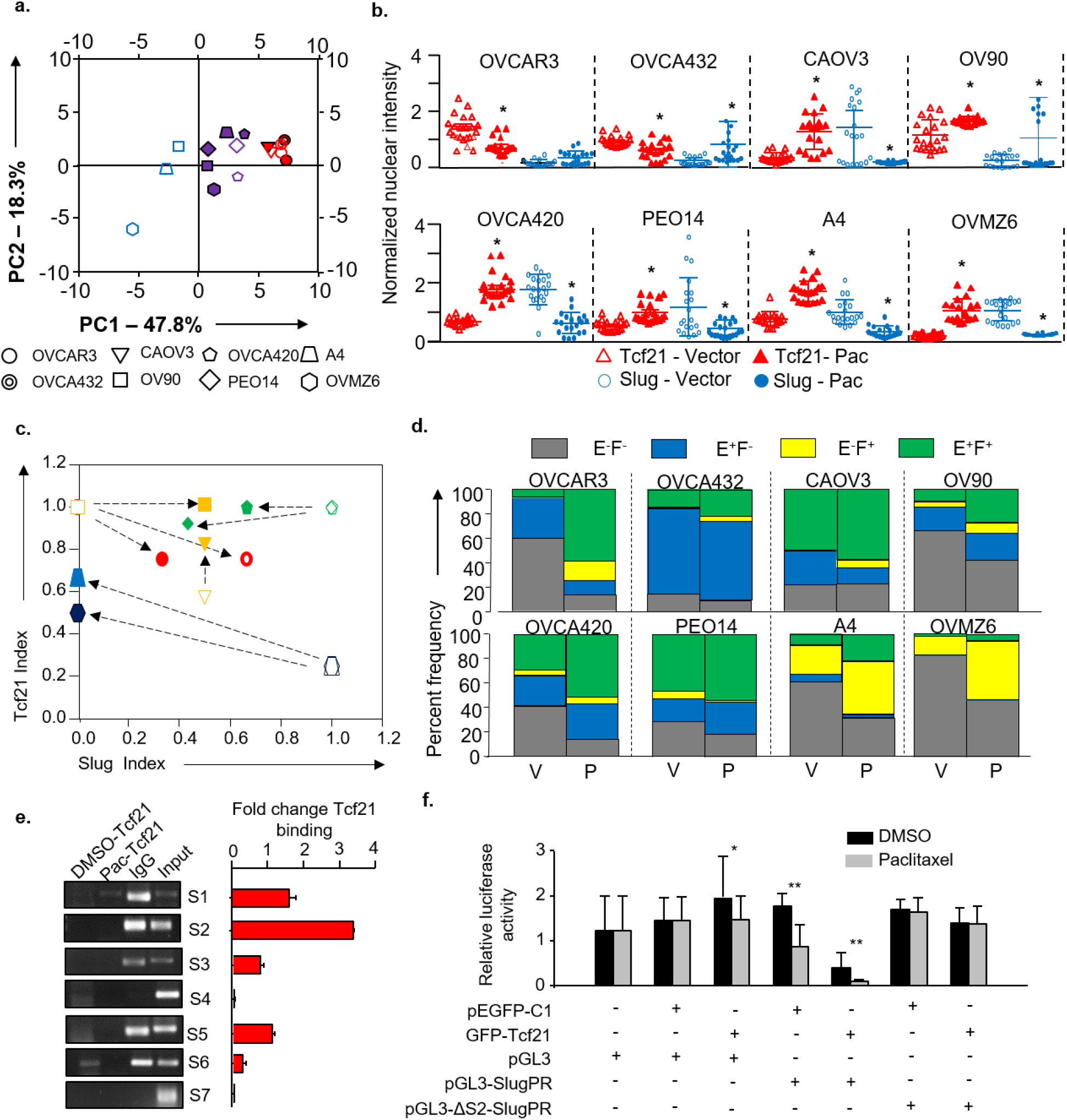
Class-switching in HGSC correlates with therapy induced Slug repression via Tcf21 a. PC analysis used to project migratory modes of the different HGSC cell lines following paclitaxel exposure, PC1 and PC2 as described earlier; empty and filled shapes correspond to 0.1% DMSO and paclitaxel exposure respectively in subsequent figures; b. Representative graphs depicting frequency and intensity of Tcf21 and Slug positive nuclei in HGSC cell lines, data depicts fluorescence intensities measured across five microscopic fields following exposure to 0.1% DMSO (vector control) and paclitaxel (Pac); c. Scatter plots of Tcf21_Index_ vs Slug_Index_ distribution for HGSC cell lines following exposure to paclitaxel; d. Graphical representation of cell population distribution based on flow cytometry profiling for Cdh1 (E) and FAP (F) levels across the phenotypic spectrum in cells exposed to 0.1% DMSO (vector control–V) or paclitaxel (P) at IC50 doses; e. ChIP-PCR analysis depicting enrichment of Slug promoter regions in Tcf21 pulldown following paclitaxel exposure of A4 cells. IgG and DMSO used as isotype and vector control respectively; f. Reporter assays to determine effects of Tcf21 on wild type Slug promoter in A4 cells, luciferase activity normalized to untransfected cells, pEGFP: empty GFP vector, GFP-Tcf21:Tcf21-pEGFPC1 recombinant vector, pGL3:native luciferase vector, pGL3-SlugPR:pGL3 containing Slug promoter, pGL3-ΔS2-SlugPR: pGL3 containing Slug promoter with deleted S2 E-box. All data are representative of experiments performed in triplicate and are depicted as mean ± SEM. *p<0.05, **p<0.01, ***p<0.001.

Application of the scores to the spectrum associated high Tcf21, low Slug indices with E and iE while an inverse profile represented the iM and M phenotype; E-M cells were depicted by high Tcf21and Slug scores. Pacliaxel exposure induced an iE phenotype in E and iE cell lines, a distinct albeit variable shift towards the E phenotype was evident in E-M, iM and M cells (Fig.6c); this was further supported by paclitaxel induced enrichment of Ecadh^+^FAP^+^ populations across the panel (Fig.6d).To assign a mechanistic understanding to therapy-induced class-switching, we probed occupancy of Tcf21 at the Slug promoter as a read-out of shift towards the E phenotype in paclitaxel treated A4 cells. Affinity of Tcf21 for the S2, S3 E-boxes in *SNAI2* promoter was not only identified following paclitaxel treatment, but also deterred in cells expressing an altered promoter in which S2 E-box was deleted. This was further complemented by negligent luciferase activity in Tcf21 over-expressing A4 paclitaxel treated cells (Figs.6e,6f); together identifying a Tcf21-driven phenotypic switch.

## Discussion

Contrasting views of the cell-of-origin of ovarian cancer being either ovarian surface or FT epithelium emerge from their anatomical proximity with sites of perturbed epithelial integrity following ovulation and wound healing^16,24,25^. The present study, restricted to the most aggressive subtype *viz*. HGSC, identifies Tcf21-Slug cross-talk in maintenance of intrinsic cellular states and tumor subtypes, wherein Tcf21-mediated Slug repression emerged as a feature of the epithelial state (Fig.7a). Rigidity of epithelial and mesenchymal states was challenged by deriving intermediates (iE,E-M,iM) and correlating phenotypes with discrete modes of migration despite comparable invasion. The latter is thus underscored as a universal feature in HGSC and uncoupled from EMT^26^. Differential migration recapitulates events at the tumor edge^27,28^, wherein destabilization of the tissue architecture by a degradative secretome and altered mileu may promote EMT, while exertion of tensile forces by adherens junction-linked acto-myosin complexes may induce CCM allowing invasion/deposition of tumor cells in the peritoneum^29–33^. Exploring physiologically relevant signaling dynamics during ovulation^34–36^, we identified BMP7-mediated induction of nuclear Tcf21, reduced migration and epithelial features, while TGF-β triggers Slug expression, EMT/aCCM and mesenchymal features. These findings may be extrapolated to derive similarities between EMT-aCCM cooperation and OSE wound healing, while maintainance of the rigid disparate epithelial state during pCCM could be akin to the FT-p53 signature-associated precursor lesions^37,38^. Adverse signaling provided by serum withdrawal or paclitaxel exposure promoted the epithelial state and CCM (Fig.7b).

**Fig7.**
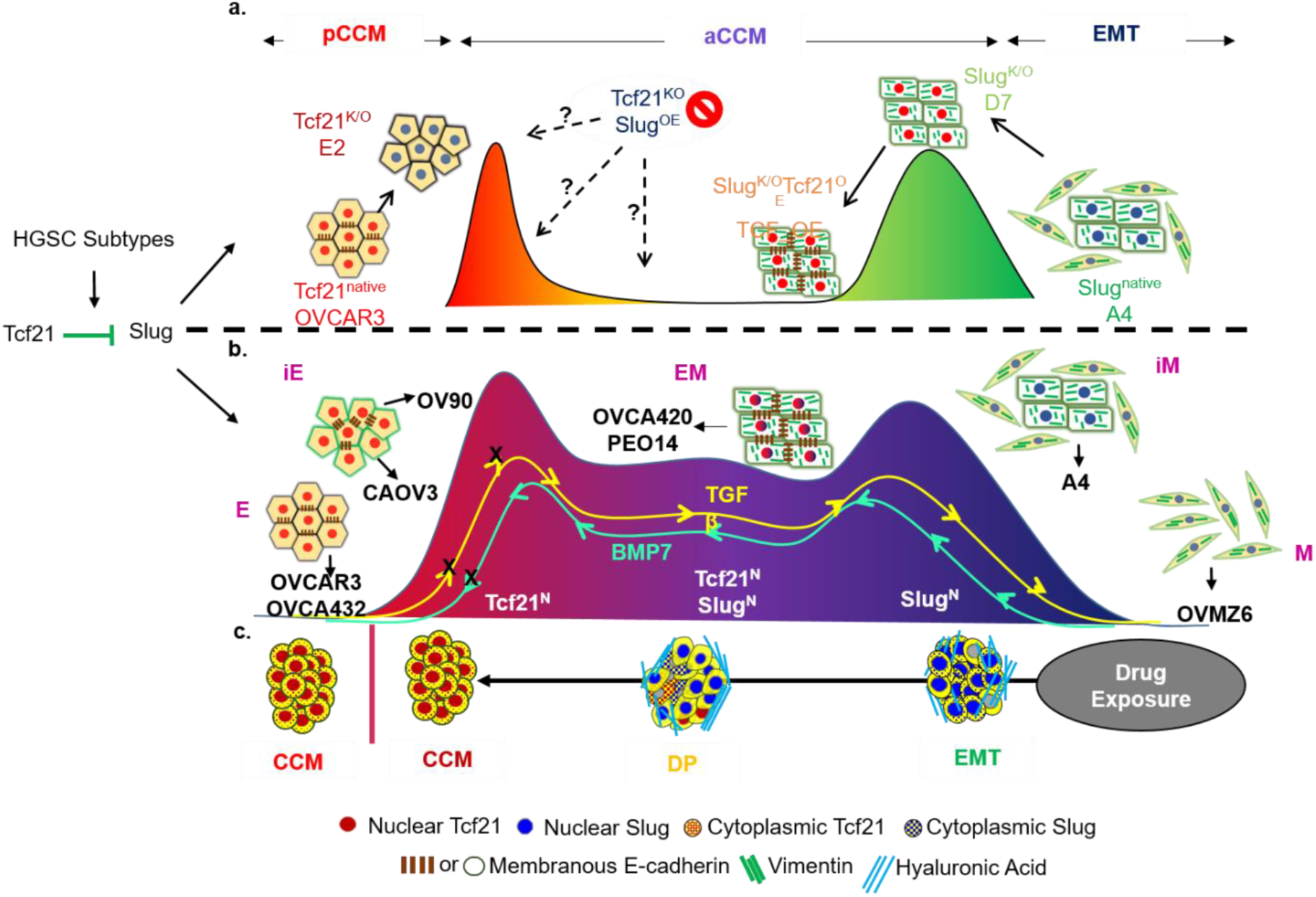
Schematic illustration depicting governance of cellular states as a function of Tcf21-Slug balance. a. Predictive derivation of Tcf21 mediated Slug repression from HGSC subtypes validates across an artificial phenotypic spectrum. K/O and OE of respective TFs result in partial phenotypic switches wherein up-regulation of Slug by Tcf21 K/O is evident. Subtleties of phenotype maintenance manifest as intermediate states observed in the Slug^K/O^Tcf21^OE^ system, while rigidity of epithelial cells is evident from lack of mesenchymal traits in Slug^K/O^ and extensive cell death of Tcf21^K/O^Slug^OE^ system. The artificial spectrum associates with distinct migratory phenotypes – pCCM, aCCM and EMT as depicted. b. Stable state spectrum identified across a panel of HGSC cell lines strengthens Tcf21-Slug mediated maintenance of cellular states identification of discrete intermediates that associate with distinct migratory phenotypes. Plasticity of intermediates is evident from transitions observed in response to positive and negative growth regulators while minimal effects are observed at epithelial end of the spectrum. c. Distinct correlations of *in vitro* phenotypes with *in situ* tumor subtypes associates therapy induced class-switching with phenotypic plasticity as a manifestation of Tcf21 mediated Slug inhibition.

*In situ* scoring of Tcf21, E-cadherin, Parp1, Slug, Anxa2 and HA emphasized association of sub-cellular localization of proteins with altered biological and functional annotations as recently ascribed^39,40^. Dominance of EMT-class in xenografts and DP-class in TMAs might reflect on cell line- or technique-associated limitations^41–43^. Importantly, stratification defined the heterogeneous in terms of intermediate phenotypes. Class-switching in response to altered niche(s) during disease progression or chemotherapy was also captured ^44–47^. These observations may align HGSC heterogeneity with distinct cells of origin by suggesting CCM- and EMT-class tumors as molecularly distinct diseases that present class-specific therapeutic targets, besides improving our understanding of cellular and biological process in the DP-class (Fig.7c;^48–54^. In conclusion, our study highlights two facets of drug resistance wherein intermediate cell states dynamically generate tumor heterogeneity, while slow-cycling epithelial cells are likely to be more resilient over other phenotypes.

## Acknowledgements

The authors acknowledge inputs in histological assessment and discussions with Dr. Avinash Pradhan, KEM, Pune, India; Statistical analysis was performed by Dr. Aditi Deshpande, Amaze Stat, Pune, India; Ms. Tejaswini Deshpande performed bioinformatics analyses for generation of the TF network. We express our gratitude to Dr. S. Mok (M. D. Anderson Cancer Center, Texas, USA) and Prof. Viktor Magdalen (Klinische Forschergruppe der Frauenklinik der TU München) for OV90 and OVMZ6 cell lines. Matlab_r2013b for analysis of live cell imaging, was provided by R.M. Deshmukh (National Centre for Biological Sciences, Bengaluru, India); Dr. A. Karthick (NCCS, Pune, India) and V.K. Vittal (Indian Institute of Science Education and Research, Pune, India) provided expertise in live image processing. This work was supported by funding to SAB from NCCS, Pune (Intramural), and Department of Biotechnology, Government of India, New Delhi (Extramural grants BT/IN/FINNISH/25/SB/2009, BT/Indo-Aus/06/03/2011). Research fellowships were availed as follows - SSV, SCK from Council of Scientific and Industrial Research, New Delhi, India; MMM from University Grants Commission, New Delhi India; BK from Department of Biotechnology, New Delhi, India, AA from NCCS, Pune, India.

## Materials and Methods

### Sample collection and preparation

The human study protocols followed all relevant ethical regulations in accordance with the declaration of Helsinki principles. The study was approved as a retrospective study by the Institutional Review Boards of National Centre for Cell Science, Armed Forces Medical College, Tata Medical Centre, Jehangir Hospital, Command Hospital and Inlaks & Budhrani Hospital. Tissue collected following surgery was formalin fixed and paraffin embedded (FFPE) using routine methods after obtaining informed consent from all patients. In all, 97 primary high grade serous ovarian adenocarcinoma cases were selected from Armed Forces Medical College (AFMC,Pune,India; 2008-2015), Tata Medical Centre (TMC,Kolkata,India; 2013-2014), Jehangir Hospital (Pune,India; 2003-2005), Command Hospital (Pune,India; 2010-11) and Inlaks & Budhrani Hospital (Morbai Naraindas Budrani Cancer Institute,Pune,India; 2013-2015).

### Animal Studies

Animal experimentation was in accordance with the rules and regulations of National Centre for Cell Science (NCCS) Institutional Animal Ethics Committee. Xenografts were raised as described earlier^16^. In brief, 2.5×10^6^ cells of cell lines OVCAR3, OVMZ6, A4, PEO14 and CAOV3 were injected subcutaneously in 3-4 week old Non-Obese Diabetic/Severe Combined Immunodeficient (NOD/SCID) mice. Animals were maintained under pathogen-free conditions and assessed every 2 days until the tumor diameter was ~1cm, whereupon animals were humanely euthanized for harvesting tumors.

### Cell culture, transfection and treatments

#### Cell Line Maintenance

Cells were cultured as per the ATCC guidelines. Ovarian cancer cell lines used in the study were all of the high grade serous histology. OVCAR3, OVCA432, CAOV3, OV90, OVCA420 and PEO14 cells were grown in RPMI 1640 supplemented with 10% fetal bovine serum. A4 cells were cultured in MEM supplemented with 5% fetal bovine serum and 1% non-essential amino acids; DMEM supplemented with 5% serum, 100μM asparagine and 100μM arginine was used for culturing OVMZ6 cells. All cell lines were maintained in a humidified incubator at 37°C under a 5% CO_2_ atmosphere. Cell line were authenticated by STR profiling and comparison with ATCC and EACC databases

#### Generation of TF derivative clones

For generation of knockouts via genomic editing OVCAR3 and A4 cells were transfected with 500ng of commercially available CrispR_GFP vectors against Tcf21 and Slug (Sigma) respectively using Lipofectamine 3000 (Invitrogen) as per the manufacturer’s protocol. Post transfection, cells were incubated at 30°C for 48 hours to ensure optimal CrispR activity. Single cell sorting of GFP positive cells was performed with BD Aria II SORP into 96 well plates. Sorted cells were allowed to grow for clone establishment and subjected to molecular (qPCR, immunoblotting, immunofluorescence staining) and functional assays (in vitro wound closure) for validation. Successful knockouts (E2 – Tcf21 knockout, D7 – Slug knockout) obtained from CrispR transfections, were further transfected with GFP vectors for Slug and Tcf21 respectively, sorted to obtain single cell clones and validated as previously described. Tcf21^K/O^Slug^OE^ clones could not be established owing to rapid apoptosis.

#### Treatments

Cells were treated as follows prior to RNA extraction, immunofluorescence staining or flow cytometry analysis:

Serum Deprivation – Cells were allowed to grow for 24 hours post seeding, following which spent medium for each cell line was replaced with equal volume of serum free media as indicated for each cell line.

Growth Factor Exposure – 24 hours following serum deprivation, cell lines were exposed to either BMP7 (10ng/mL), TGFβ (10ng/mL) or a combination of both (Invitrogen).

Paclitaxel Treatment – Cells were allowed to grow for 24 hours post seeding, after which spent medium for each cell line was replaced with equal volume of complete medium containing IC50 concentrations of paclitaxel (Sigma) optimized for each cell line (IC50 values for each cell line are available on request).

### Derivation of TF network, correlation and expression analysis of TFs and phenotypic markers in TCGA-HGSC dataset

#### Derivation of TF network

As previously reported^16^, weighted gene correlation network analysis (WGCNA) identified differential expression of gene modules across discrete molecular subclasses. Class1 (CCM) and Class2 (EMT) exhibited differential expression of 3 modules each which were examined to identify regulatory genes. Briefly, top 25 genes from each of the differentially expressed modules were reviewed in literature to identify transcription factors. From the 13 TFs identified 2 immunomodulatory TFs were eliminated from the study due to technical limitations in assessing these genes in HGSC cell lines and immunocompromised mouse models. Transcription factor binding site analysis involved defining the promoter region for each TF (−3kn to +1kb from transcription start site) and assessing regulatory elements for other TFs in the same. Based on identification of respective binding sites and review of expression data for the TFs a directional TF network was predicted.

#### Correlation and Expression Analysis of TFs and phenotypic markers

FPKM values available for gene expression data on the TCGA-GDC (Genomic Data Commons) Portal for 379 HGSC tumor samples were downloaded and utilized to extract data for the 11 TFs identified in the network and epithelial, mesenchymal markers was extracted from the TCGA-HGSC dataset. List of epithelial and mesenchymal phenotypic markers was obtained from an exhaustive PubMed review involving the search terms, ‘ovary AND epithelial OR mesenchymal’. Correlation co-efficient obtained for the phenotypic markers was subjected to a filter of R^2^<-0.2 and >0.2 across the dataset to eliminate poorly correlating genes. Data representation was performed using the MeV software.

For depicting expression profiles of E and M TFs in the HGSC dataset, FPKM data was log_2_ normalized, median centered and represented as a heat-map.

### RNA extraction, cDNA synthesis, RT-PCR, qPCR and PCA analysis

#### RNA extraction, cDNA synthesis, RT-PCR, qPCR

Trizol based total RNA extraction was achieved from 80% confluent cultures of cell lines following respective treatments. Briefly, Trizol suspensions were subjected to phase separation with chloroform, followed by collection of the aqueous layer. RNA precipitation by 100% iso-propanol was followed by two washes with 70% ethanol prepared in DEPC treated water. cDNA synthesis was performed using SuperScript™ First-Strand Synthesis System (Invitrogen) as per manufacturer’s protocol. RT-PCR and qPCR reactions using gene-specific primers were performed in triplicate. qPCR reactions were performed using Applied Biosystems Step One Plus instrument in 96- well plate format using SYBR Green PCR Master Mix (Applied Biosystems). Changes in threshold cycle (Ct) values between the gene of interest and endogenous control was calculated as: ΔCt = Ct (test gene) – Ct (GAPDH); relative expression was calculated as: 2^-(Δ Ct)^. Fold change in expression was calculated as: ΔΔCt = ΔCt (test gene - treatment) – ΔCt (test gene – control). GAPDH expression was used as an endogenous control; non-template controls were used to eliminate background readings contributed by probable contaminants in the reaction mixture. All expression data was log_2_ normalized for better representation. Primer sequences have been provided in the key reagents and resources section.

#### Principal Component Analysis (PCA) of qPCR data

Fold expression change data for TFs and markers obtained from qPCR following specific treatments were analysed in matlab2013b (Mathworks) with an open source code available for principal component analysis. First two components capturing maximum variance of the data were represented using SigmaPlot 10.0 software (SPSS).

### Immunoblotting

Immunoblotting protocols were followed as previously described^55^. Briefly, cell lysates were prepared in RIPA buffer (50mM Tris – pH 7.4, 150mM NaCl, 2mM EDTA – pH 8.0, 1% NP-40, 0.1% SDS) containing 2mM PMSF and 1X protease inhibitor cocktail (Roche). Proteins estimation was performed as per manufacturer’s instructions with the DC protein assay kit (Bio-Rad). 50 μg protein denatured with 5X SDS loading buffer (10%SDS, 10mM β-mercaptoethanol, 20% glycerol, 0.2M Tris-Cl pH 6.8, 0.05% bromophenol blue), were resolved by electrophoresis on 12.5% SDS–PAGE gels. Resolved proteins were transferred onto a PVDF membrane (Amersham Pharmacia Biotech), blocked for 1 hour in 5% bovine serum albumin in TBS containing 0.1% Tween 20, and then incubated over-night with specific primary antibodies at 4°C. Membranes were then washed and incubated with specific horseradish peroxidase (HRP) linked secondary antibodies (dilution 1:5000) for 1 hour at RT. Signals were detected using SuperSignal West Pico Chemiluminescent Substrate (Pierce) as per the manufacturer’s instructions. The primary-secondary antibody complex was dissociated from the membranes, via stripping with 100mM β-mercaptoethanol, 2% SDS and 62.5mM Tris–HCl (pH 6.8) for 20 min at 50°C, followed by immunoblotting as mentioned above. A comprehensive list of antibodies used is provided in the key reagents and resources section.

### Immunostaining, image acquisition and quantification of signal intensity

#### Immunostaining and image acquisition

Cells grown on coverslips were fixed with 4% paraformaldehyde for 10 min at 4 °C post specific treatments. Fixed cells were then permeabilized with chilled methanol for 10 min, blocked with 5% BSA for 1 h and incubated with specific primary antibody for 1 hour. Cells were then incubated with secondary antibody for 20 minutes, and nuclei were counter-stained with Hoechst 33432 (Thermo Fisher) for 10 minutes. Intermittent washes were provided with 1X phosphate buffered saline (PBS). Images were acquired on an inverted laser scanning confocal fluorescence LPSS5 microscope (Leica Microsystems, Wetzlar, Germany; 63× oil immersion objective). A comprehensive list of antibodies has been provided in the key reagents and resources section

#### Quantification of signal intensity

Quantification of signal intensities for transcription factors was performed with the LAS-AF software (Leica Microsystems). Briefly, ROIs were defined around nuclear margins of each cell followed by extraction of signal intensities for Hoechst and respective fluorescent channels. Signal intensity for specific transcription factors was determined as a signal ratio of TF-associated fluorescence/Hoechst. Quantification was performed for a minimum of 50 nuclei for each sample.

### Boyden chamber invasion assay

Trans-well invasion assays were performed, as previously described (Kurrey *et al*., 2009). Briefly, a suspension of 2.5× 10^4^ cells in serum free medium was seeded on 5% Matrigel (BD Bio-Science) coated polycarbonate membranes (8 μm pore size) in trans-well culture chambers. 500 μl medium (serum free or containing 5% serum) was filled in the lower compartment of the chamber. After 48 h of incubation at 37°C & 5% CO_2_ atmosphere the membrane and matrigel were fixed with 4% paraformaldehyde, stained with 0.01% crystal violet and counted in 10 different fields for invading cells under an Olympus IX71 microscope at 10X magnification.

### *In vitro* wound healing assay, live cell imaging, processing and quantification of data, PCA for migration

#### In vitro wound healing assay

Wound healing assay was performed as per previous protocols ^16^.1 × 10^5^ cells/well were seeded in 24-well plates and incubated at 37 °C over-night to achieve 90% confluency. Wound was inflicted on the monolayer with a pipette tip, washed with 1XPBS twice, followed by addition of appropriate volume of serum free medium. Under specific experimental conditions media were additives administered to the media included 5% serum, TGFb (10ng/ml), BMP7 (10ng/ml) or a combination of both growth factors. The wound was allowed to heal and was monitored upto 144h. Images were captured on Olympus IX71 microscope and analyzed by the T-Scratch Software (Tobias Gebäck and Martin Schulz).

#### Live Cell Imaging

Images were acquired for 30 hours, following infliction of wound on a cell monolayer, on an inverted laser scanning confocal fluorescence LPSS5 microscope maintained at 37°C, 5% CO_2_ atmosphere (Leica Microsystems). In case of specific treatments, additives were added as previously described. For live cell imaging of paclitaxel treated cells, drug treatment for 48 hours was followed by harvesting of residual cells and seeding for wound healing assay. In the absence of additives live cell imaging was performed in serum free media in the presence of 20ng/mL of mitomycin ‘C’ (Sigma),

#### Processing and quantification of data

Images captured for live cell imaging were exported from the LAS AF software with.avi extensions. For elimination of shadow effects in images, due to non-specific anomalies in the plate architecture,.avi files were opened in Fiji software, duplicated, inverted and averaged with original image using the Image Calculator. Contrast adjustments were made as per requirement. Processed files were subjected to thresholding to obtain binary images which were eventually processed through MTrack2 plugin to obtain X and Y co-ordinates. X and Y co-ordinates over a period of 16 hours was used to calculate mean displacement and velocity for each cell. Furthermore, binary images were processed through a TrackMate plugin to determine the mean number of nearest neighbours.

#### PCA for migration

Mean displacement, velocity and number of nearest neighbours were analysed in matlab2013b (Mathworks) with an open source code available for principal component analysis. First two components capturing maximum variance of the data were represented using SigmaPlot 10.0 software (SPSS).

### Suspension culture assessment

Suspension cultures for HGSC cell lines were generated as described by earlier studies^56^. Briefly, 5 × 10^4^ cells/well were seeded in ultra-low attachment plate (Corning) in serum free medium and incubated at 37 °C for 14 days. The suspension culture morphology was imaged on Olympus IX71 microscope.

### Clonogenecity assay

Clonogenecity assay preformed as per previous description^56^. Briefly, 1 × 10^3^ cells/well of each cell line were plated in 96-well plates followed by incubation at 37 °C for 72 hours; adherent colonies were washed twice with 1XPBS, fixed with 4% paraformaldehyde, stained with 0.05% crystal violet, and imaged with Olympus IX71 microscope. Colony counts were obtained with the ImageJ software

### FACS sorting and flow cytometry

For detection of specific markers by flow cytometry, cells were grown to 80% confluency prior to harvesting with 0.5% Trypsin-EDTA. Cells were fixed with 4% paraformaldehyde followed by pearmeabilization with ice cold 50% methanol for 10 minutes. Staining for membrane associated markers proceeded without permeabilization. Non-specific antibody binding was blocked with 5% BSA, followed by incubation with respective primary and secondary antibodies. Acquisition of stained cells was performed using BD FACS Canto (BD Biosciences) and analyzed with the BD FACS Diva software version 6.0 (BD Biosciences) and FlowJo (). A comprehensive list of antibodies is provided in Star Methods

### Cloning, expression and purification of GST-Tcf21 protein

Human Tcf21 cDNA was amplified from A4 cells, cloned into pGEX-6P-1 GST vector and transformed into Escherichia coli BL21 codonPlus (Agilent Technologies). Cells grown overnight in LB medium (100 μg/ml ampicillin and 34 μg/ml chloramphenicol) at 37 °C were sub-cultured in Terrific Broth medium (100 μg/ml ampicillin + 34 μg/ml Chloramphenicol) at 37 °C to an optical density of 0.6–0.8 at 600 nm. Protein expression was induced by 0.5mMisopropyl β-D-1-thiogalactopyranoside (IPTG) at 16 h at 22 °C. Cells were sonicated in lysis buffer [50 mM Tris–HCl, 300 mM NaCl, 5 mM DTT, 1 mM PMSF and Protease inhibitor cocktail, pH 7.5] and equilibrated with Glutathione sepharose (Amersham Biosciences). GST-tagged Tcf21 fusion proteins were purified by affinity chromatography under native conditions and concentration estimated by DC protein assay kit (Bio-Rad), using bovine serum albumin as protein standard. GST-Tcf21 purification was validated by immunoblotting for Tcf21 and GST as described above.

### Electrophoretic mobility shift assay (EMSA)

30-bp double-stranded oligonucleotide probes were designed and synthesized around each E-box element in the putative human Slug promoter regions. Synthesized oligonucleotides annealed in NEB2 (New England Biolabs) buffer were used in EMSA. Binding was assayed with 50 ng purified GST-Tcf21 and GST protein, 5μg reannealed oligonucleotide probe and buffer (100 mM HEPES (pH 7.5), 2.5 mM DTT, 50% glycerol, 25 mM MgCl2, 0.25 mM ZnSO4) at room temperature for 20 min. EMSAs were run at 4°C on 6% polyacrylamide gel in 0.5× tris–boric acid–EDTA (TBE) buffer, electrophoresed at 100 V for 60 min and visualized by ethidium bromide staining. 30-bp probes lacking E-box sequences as controls. A comprehensive list of oligonucleotides used is provided in key reagents and resources table.

### Chromatin Immunoprecipitation Assay and ChlP-PCR

ChIP was performed as described earlier (Kurrey *et al*., 2009); ChIPped DNA amplified using the GenomePlex Complete Whole Genome amplification kit as per manufacturer’s instructions (Sigma-Aldrich). Specific primers were used to amplify E-boxes within promoter regions of genes, amplified products run on 1.5% agarose gels, and band intensities measured by densitometric analysis using Genetool3.6 (Syngene). A comprehensive list of primers is provided in key reagents and resources table.

### Model generation

A mathematical model to integrate expression data with reported Tcf21, Slug interactions was developed^55,57^. The model formulation for is given as:

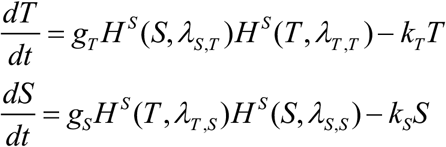

where T and S denote TCF21 and SLUG respectively. *g_T_* and *g*_*S*_ are respective production rates for TCF21 and SLUG, and *k_T_* and *k_S_* are their respective degradation rates. Shifted Hill functions, denoting the effect of X on Y, are defined as *H^s^* (*X,λ*_*X,Y*_) = *H^−^* (*X*) + *λ_X,Y_H*^+^ (*X*) where *H*^−^(*X*) is the inhibitory Hill function, *H*^+^ (*X*) = 1 – *H*^−^ (*X*) is the excitatory Hill function, and *λ_X,Y_* denote the equivalent of fold-change in production of Y due to X^58^. Inhibitory Hill function is defined as: 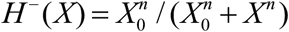 where *X*_0_ is the half-maximal threshold, and *n* is effective cooperativity.

Degradation rates for TCF21 and SLUG (represented in per unit hour; assuming first-order kinetics) are estimated from experimental data on their half-lives^59,60^. Production rate (represented in 1000 molecules per hour) is estimated, based on total number of protein molecules per cell, as reported for signaling molecules (~100,000; Milo *et al*., 2010). Fold-change for effect of Slug on Tcf21 and vice-versa has been estimated from our results and those for self-activation have been gathered from existing data. The parameters used here are:

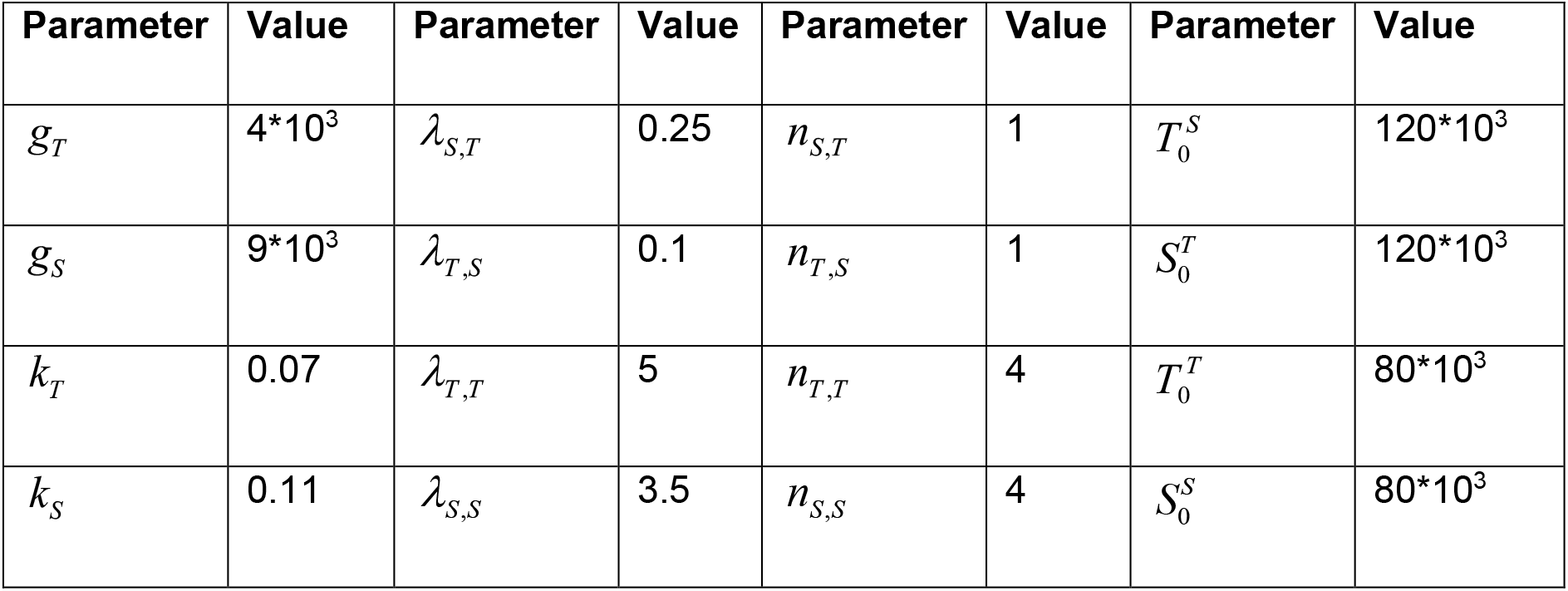

### Luciferase Constructs and Luciferase assay

Luciferase constructs were for E-cadherin and Slug wild type promoters along with deletion mutants ΔS1–S2 were previously generated in the pGL3 luciferase plasmid^55^. All constructs were verified by sequencing. 0.5 × 10^5^ OVCAR3, A4, and derivatives were cells were transfected with 0.8 μg Luciferase reporter plasmids control and Slug promoter containing plasmids in 24-well plates using Lipofectamine 3000 (Invitrogen). Renilla luciferase was added (10 ng) to each transfection as control. Luciferase activity was measured using a Dual Luciferase assay kit (Promega) either post transfection with pEGFP-Tcf21 and/or post paclitaxel treatment. Empty pEGFP and pGL3 vectors were used as controls.

### Immunohistochemical (IHC) and Histochemical staining (HC)

IHC and HC based read-outs in xenografts (one cell line per phenotype – E: OVCAR3, iE: CAOV3, E-M: PEO14, iM: A4, M: OVMZ6) were selected for assessment. IHC and HC was performed in 5μm sections of FFPE blocks fixed by drying at 60°C for at least 1h in oven using standard protocol, deparaffinized in xylene and hydrated in ethanol-distilled water gradient. Heat-induced epitope retrieval (HIER) was carried out for 30min at pH=9 / pH=6. For peroxidase inactivation sections were incubated in 3% H_2_O_2_ for 30min, followed by 1x Blocking Solution for 10min (Biogenex) and overnight incubation in primary antibodies. Sections were washed and incubated with specific secondary antibodies for 1hr, color developed with DAB (Thermo Pierce); hematoxylin used as a counter stain. Sections were further dehydrated and mounted in DPX (Qualigens). Negative controls were prepared in the absence of primary antibody. For HC-based HA detection, test sections were exposed to freshly prepared hyaluronidase (Sigma-Aldrich); control slides were incubated in phosphate buffer for 1hr at 37°C. Sections were washed in running water for 10min and stained with Alcian blue for 30min (Fluka), counter-stained with Nuclear Fast Red Solution for 2min (Sigma-Aldrich), dehydrated and mounted in DPX (Qualigens). Positive experimental controls included testis (TCF21, PARP1), liver (E-cadherin), lymphocytes (Slug), gall bladder (ANXA2) and small intestine (hyaluronan); negative controls included heart (TCF21, E-cadherin, Slug, ANXA2) and mucosa of small intestine (PARP1). IHC and HC methods were standardized for each marker as Standard Operating Procedures (SOPs). Reproducible SOPs were established to address pre-analytic (slide coating, tissue selection, fixation, processing), analytic (clone and antibody selection, buffers and instruments for antigen retrieval, antibody/enzyme concentration, duration of each step, etc.) and post-analytic parameters (interpretation, analysis and reporting of expression in the reference and control tissues). Slides for human tissues, xenografts and TMA were reviewed independently (AS,RDD,ANJ,DM,SCK); a consensus was reached to establish tissues for reference score. Human patient cohort slides were reviewed by AS,RD,AJ,SCK.

### Scoring guidelines

Specific scoring guidelines for each marker included –

i. *Score for Marker Frequency (S_Freq_)* – percentage expression in total tumor cells of tissue section on a scale of 0 - 3 (0: absent, 1: 1-10%, 2: 11-50% and 3: ≥ 51% marker positive),

a. TCF21: cardiac myocytes, ovarian stromal cells, germinal cell of testis represented S_Freq_ 0, 1 and 3 respectively; S_Freq_ = 2 could not be identified in healthy tissues.
b. E-cadherin: cardiac myocytes, liver hepatocytes, prostate epithelial cells represented S_Freq_ 0, 2 and 3 respectively; healthy tissues representing S_Freq_ = 1 could not be identified.
c. PARP1: mucosa of small intestine, cardiac myocytes, germinal basal cells of testis represented S_Freq_ as 0, 1 and 3 respectively; healthy tissues representing S_Freq_ = 2 could not be identified.
d. Slug: cardiac myocytes, smooth muscles of appendix, lymphocytes of small intestine represented S_Freq_ 0, 1 and 2 respectively; healthy tissues representing S_Freq_ = 3 could not be identified.
e. HA: cartilage and sub-mucosa of small intestine represented SFreq as 2 and 3 respectively; healthy tissues representing S_Freq_ = 0 or 1 could not be identified.
f. ANXA2: cardiac myocytes, somatic muscle of small intestine, epithelial cells of gall bladder represented S_Freq_ 0, 1 and 3 respectively; healthy tissues representing S_Freq_ = 2 could not be identified.
ii. *Score for Marker Intensity (S_Int_)* – intensity of brown stain for IHC and blue for HC in positively stained tissue sections. A scale of 0 – 3 was established, 0: absent, 1: weak, 2: moderate and 3: strong intensity of marker positive cells,

a. TCF21: cardiac myocytes, ovarian stromal cells, germinal basal cells of testis represented S_Int_ 0, 1 and 2 respectively; S_Int_ = 3 could not be identified in healthy tissues.
b. E-cadherin: cardiac myocytes, epithelial cells of small intestine, epithelial cells of prostate represented S_Int_ 0, 2 and 3 respectively; healthy tissues representing S_Int_ = 1 could not be identified.
c. PARP1: mucosa of small intestine, cardiac myocytes, germinal basal cells of testis represented S_Int_ 0, 1 and 2 respectively; healthy tissues representing S_Int_ = 3 could not be identified.
d. Slug: cardiac myocytes, smooth muscle of appendix, lymphocytes of small intestine represented S_Int_ 0, 1 and 2 respectively; healthy tissues representing S_Int_ = 3 could not be identified.
e. HA: Intensity for hyaluronan was measured as blue color intensity developed by Alcian blue in comparison to hyaluronidase digested tissue section. Sub-mucosa of small 6 intestine and cartilage tissues represented S_Int_ 1 and 2 respectively; healthy tissues representing S_Int_ = 0 or 3 could not be identified.
f. ANXA2: cardiac myocytes and epithelial cells of gall bladder represented S_Int_ 0 and 2 respectively; healthy tissues representing S_Int_ = 1 or 3 could not be identified.
iii. *Score for Marker Localization (S_Loc_)* – representing sub-cellular location of marker in the tissue section on a scale of 0 to 2, 0: Absent, 1: mislocalized (cellular localization does not correspond to known functionality, for example, cytoplasmic location for TCF21, PARP1, Slug, E-cadherin, ANXA2 or HA), 2: normal localization (for example, nuclear expression of TCF21, PARP1 or Slug, membrane for E-cadherin, membrane or cytoplasmic for ANXA2 and extracellular expression of HA.

a. TCF21: cardiac myocytes, liver hepatocytes, germinal bas*al cells of testis represented S_Loc_ 0, 1 and 2 respectively.
b. E-cadherin: cardiac myocytes, prostate epithelial cells represented S_Loc_ 0 and 2 respectively; healthy tissues representing S_Loc_ = 1 could not be identified.
c. PARP1: mucosa of small intestine, germinal basal cells of testis represented S_Loc_ 0 and 2 respectively; healthy tissues representing S_Loc_ = 1 could not be identified.
d. Slug: cardiac myocytes, somatic muscle of appendix, lymphocytes of small intestine, represented S_Loc_ 0, 1 and 2 respectively.
e. HA: cartilage represented S_Loc_ of score 2; healthy tissues representing S_Loc_ = 1 could not be identified. A further consensus was reached in the pathology review to consider extracellular staining in tumor nests that is eliminated following hyaluronidase treatment as a proper localization, while distant stroma-associated HA was considered as mislocalization.
f. ANXA2: cardiac myocytes, stromal cells of gall bladder, epithelial cells of gall bladder represented S_Loc_ as 0, 1 and 2 respectively.

### Quantitative and Statistical Analysis

Pearson’s Correlation Co-efficient across the TCGA-HGSC datasets was calculated using the GraphPad Prism 6.0 software.

Significant differences in gene expression profiles across the HGSC cell line panel were calculated by One-Way ANOVA using the Sigma Stat 3.5 Software. Differences were considered significant if p<0.05.

Significant differences following treatments in gene expression profiles, invasive capabilities and wound healing properties of HGSC cell lines and derivative clones, and luciferase activity for promoter assays were calculated by paired Student’s t-test using the Sigma Stat 3.5 Software. Similarly, ChIP binding efficiencies were calculated by un-paired Student’s t-test Differences were considered significant if p<0.05.

Differences in nuclear intensities of Tcf21 and Slug were calculated by Tukey’s Test using the Sigma Stat 3.5 Software. Differences were considered significant if p<0.05.

Biomarker scores for frequency, intensity and localization collected from each observer were used to compute Biomarker and Class indices. These were compared between groups by Pearson correlation using SPSS for Windows; Student’s t-test and ANOVA were determined in Microsoft Excel 2016; p<0.05 was considered significant.

### Data Availability Statement

The authors declare that all relevant data supporting findings in this study are available within the paper and its supplementary information files. SOPs developed in this study are available from the corresponding author upon reasonable request.

